# SwinCell: a transformer-based framework for dense 3D cellular segmentation

**DOI:** 10.1101/2024.04.05.588365

**Authors:** Xiao Zhang, Zihan Lin, Liguo Wang, Yong S. Chu, Yang Yang, Xianghui Xiao, Yuewei Lin, Qun Liu

## Abstract

Segmentation of three-dimensional (3D) cellular images is fundamental for studying and understanding cell structure and function. However, 3D cellular segmentation is challenging, particularly for dense cells and tissues. This challenge arises mainly from the complex contextual information within 3D images, anisotropic properties, and the sensitivity to internal cellular structures, which often lead to miss-segmentation. In this work, we introduce SwinCell, a 3D transformer-based framework that leverages Swin-transformer for flow prediction and effectively distinguishes individual cell instances in 3D. We demonstrate the broad utility of the SwinCell in the segmentation of nuclei, colon tissue cells, and dense cultured cells. SwinCell strikes a balance between maintaining detailed local feature recognition and understanding broader contextual information. Tested extensively with both public and in-house 3D cell imaging datasets, SwinCell shows superior performance in segmenting dense cells in 3D, making it a powerful 3D segmentation tool for cellular analysis that could expedite research in cell biology and tissue engineering.

## Introduction

Cells, the fundamental units of living organisms including humans and plants, are pivotal subjects in biological research. Understanding their behavior, tissue organization, and implications in disease progression and treatment necessitates an exploration of their three-dimensional (3D) spatial context. Various 3D imaging techniques, such as 3D holotomography^1^, X-ray computed tomography^2^, confocal microscopy, cryo-electron tomography (cryo-ET)^3,4^, and volume Electron Microscopy (vEM)^5^ have been widely adopted, generating vast amounts of 3D data. Consequently, there is an urgent need for corresponding advances in image analysis algorithms to effectively analyze these 3D data.

In this context, 3D segmentation plays a crucial role. It enables accurate quantification of cell volume, shape, and intercellular spatial relationships, providing essential insights into biological processes. Particularly in areas like drug development, where the efficacy of treatments can be profoundly influenced by the 3D arrangement of cells and tissues, precise segmentation algorithms are indispensable^6,7^. Therefore, the development of advanced 3D cell segmentation algorithms is pivotal, given the intrinsic three-dimensional nature of biological tissue and cellular structures.

Recent advancements in deep-learning-based segmentation algorithms have facilitated accurate pixel-level segmentation of 2D images^8–11^. Traditional deep-learning models like U-Net^12^ and Mask-RCNN^13^ have been widely used for 2D image segmentation, offering robust performance across various imaging modalities. However, the segmentation of dense cells and tissue images, especially for 3D cellular images, remains challenging. To overcome the drawback of these 2D models, shape representations, such as gradient flows^10,14^, distance maps^9^, and star-convex polygons^15^ were introduced to improve the separations of adjacent cells. Among these algorithms, Mesmer^9^ combines Feature Pyramid Network and distance maps for dense cell segmentation. Similarly, the Cellpose model ^10,11^ predicts simulated flow gradients along each image dimension and utilizes gradient tracking to group connected cell pixels into masks. The integration of flow gradient prediction helps the Cellpose algorithm achieve the state of the art performance.

Previous efforts have been made to extend the 2D segmentation algorithms to 3D. For instance, StarDist3D extends the 2D StarDist^16^ algorithm to 3D by extending the prediction of 2D star-convex polygons to 3D polyhedra^15^. This approach, however, relies on the assumption that cells inherently possess star-convex geometries, an assumption that may not always hold. Another key challenge in using StarDist3D for adherent cell segmentation lies in its application of Non-maximum Suppression (NMS) in the post-processing step, which inadvertently suppresses valid cell representations in scenarios where cells are densely packed. As a result, NMS-based algorithms usually suffer low recall problems^17^. While Cellpose and Mesmer algorithms can be extended to 3D segmentation by predicting 2D masks of slices in each plane and stitching them together, this approach is prone to artifacts resulting from the stitching process.

The primary challenge in 3D segmentation is the model’s capability to effectively learn and interpret 3D shape representations. Accurate 3D instance segmentation requires a deep understanding of the complex shapes and contexts within 3D images. The convolutional neural networks (CNNs) employed in both Cellpose and Mesmer naturally possess restricted receptive fields, limiting their capacity to capture contextual information solely within localized regions. In contrast, Transformers^18^ operate by encoding visual representations from a series of patches and utilize self-attention mechanisms to effectively model long-range context information. Recently, the Shifted Windows (Swin) Transformer^19^ has been proposed as a general-purpose backbone that employs a hierarchical Transformer architecture, enabling localized self-attention computations with non-overlapping windows.

In this work, we developed a 3D SwinCell framework that incorporates the Swin-transformer architecture and 3D U-net to provide cohesive and unified flow predictions, achieving more accurate cell mask prediction and segmentation. The 3D SwinCell algorithm was extensively tested on multimodal datasets, including two publicly available datasets and one in-house dataset collected with Nanolive 3D holotomography ^1,2^. The 3D SwinCell algorithm outperforms the state-of-the-art Cellpose algorithm^2,3^ in the application of different cell segmentation scenarios.

## Results

### SwinCell workflow and model

The SwinCell workflow (**Fig. 1**) starts with manually labeling a subset of raw 3D images to generate training data with 3D masks, distinguishing dense cell boundaries from the background (**Fig. 1a**). The masks are then transformed into instance segmentation masks and flow representation of cells (**Fig. 1b**). The flows of the cells provide more geometric information of the cells and allow the SwinCell model to segment individual cells by considering the direction and magnitude of pixel displacements.

**Fig. 1.**
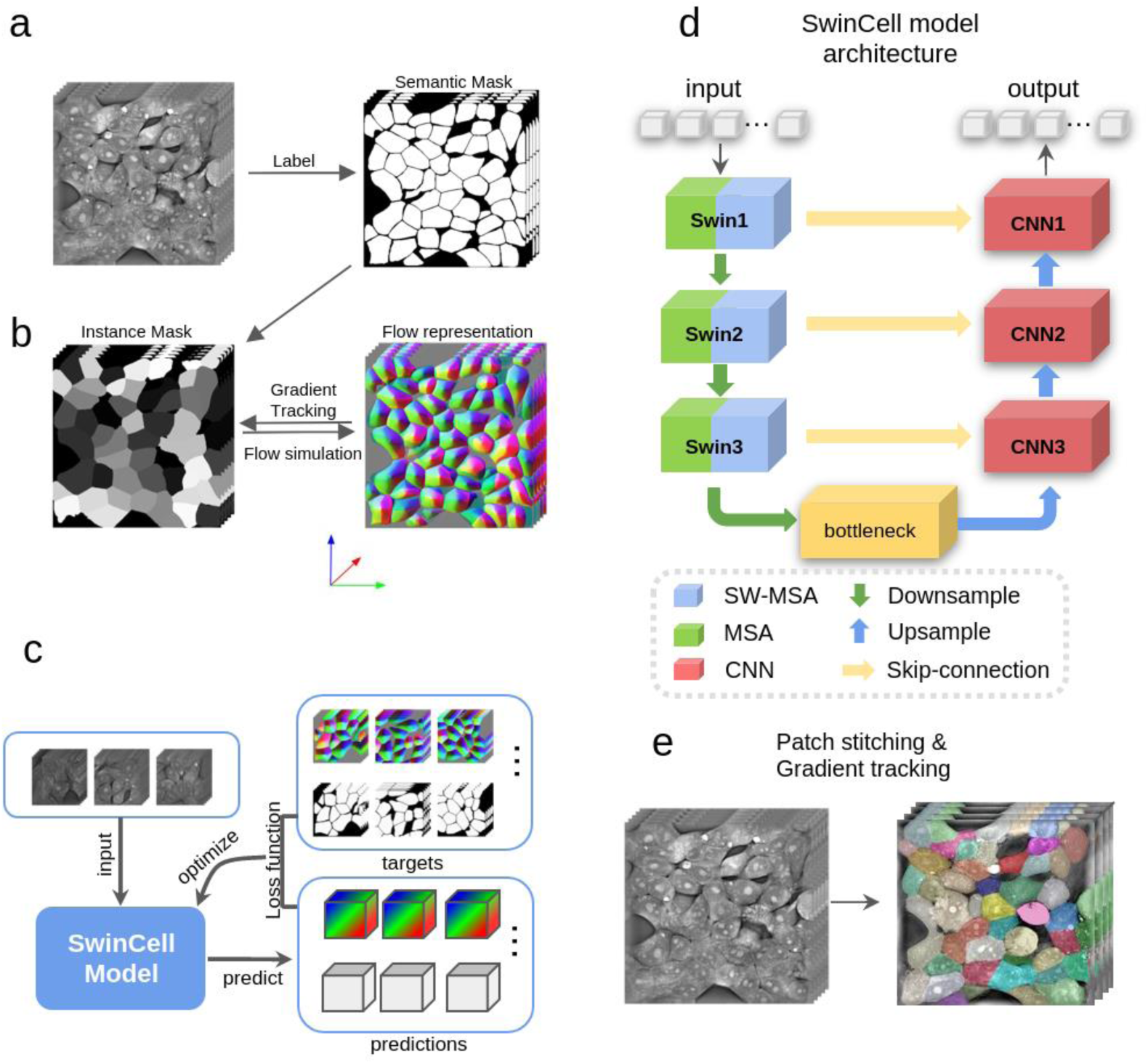
SwinCell workflow. **a,** Image labeling and annotation. Raw 3D cellular images are manually labeled by separating individual cells for subsequent processing. **b**, Label transformation and flow generation. Semantic cell labels are converted to instance labels, which are then transformed into gradient flows using the algorithm described in the Method section. The RGB components of the color-encoded flow map represent gradient flows in the Z, X, and Y directions, respectively. **c**, Model training. The training process involves randomly cropping 3D image patches (128 x 128 x 32) embedded with their corresponding labels. The model inputs are embedded raw images and outputs are four-channel predictions. The first three channels are gradient flows in Z, X, and Y directions, and the fourth channel is the cell probability map. A loss function is computed from the model predictions and the ground truth, which is then used to back-propagate errors and update the model’s parameters. **d**, Model architecture. SwinCell utilizes a U-net-style architecture with three levels. Each level in the encoder section contains a Swin-Unit, and the decoder section utilizes convolutional upsampling. **e**, Model Inference. The model takes raw images as input and produces four-channel predictions. Predicted image patches are stitched together to reconstruct the complete cellular image. Gradient tracking is then applied to convert the predicted channels into a cell instance mask.

Due to the large size of 3D raw image and label pairs, which often exceeds the memory capacity of a standard GPU, smaller image patches are randomly extracted from the original image and used as model inputs for training (**Fig. 1c**). The flow representations can either be calculated in real-time during the training process or pre-generated and stored for more efficient training in the preprocessing steps. The patches are used as inputs by the model to predict flows and cell probability maps, and loss functions are computed separately for both the model-predicted flows and the cell probability maps, compared to the ground truth. These calculated losses are then used to update the model parameters through back-propagation.

The SwinCell model utilizes a Swin-transformer to enhance flow predictions from input images. The model architecture is illustrated in **Fig. 1d**. The input to the model is 3D image patches. Similar to the traditional U-Net^12,20^ model architecture, SwinCell contains an encoder and a decoder, and the two parts are connected by skip connections and a bottleneck. The encoder has three Swin-Transformer units (**Extended Data** Fig. 1) and each unit contains a standard multi-head self-attention (MSA) and a shift-window multi-head self-attention (Swin-MSA). In the encoder part, the resolution of features is down-sampled by a factor of 2 at each stage. In the decoder part, the resolution of the features is increased by a factor of 2 at each stage using a convolutional style up-sampling layer. This multi-level design allows the model to learn image features at different resolutions. The trained SwinCell model takes input image patches, predicts segmented patches, and stitches them to the original image size with Gaussian weighted averaging (**Fig. 1e**). Gradient tracking^10,14^ is then applied to the stitched images to generate instance segmentation masks. Each cell is assigned a unique integer ID, represented by distinct colors (**Fig. 1e**).

### Benchmarking SwinCell model performance with a public Allencell (hiPS) dataset

The Allencell (hiPS) dataset ^21^ contains 200,000 single cells from over 18,000 live-cell images. The dataset includes several fluorescent protein channels that mark different cellular components, offering a three-dimensional view of cellular structures ^21,22^. In this work, the nucleus channel from the first 100 images was selected to evaluate the model performance because the nuclei are well recognized and away from each other (**Fig. 2a**). The qualitative and quantitative comparison of segmentation results between our SwinCell model and the Cellpose model are shown in **Figs. 2b-f**. We chose to use the Cellpose model for benchmark comparison because it is widely recognized and commonly used for cell segmentation.

**Fig. 2.**
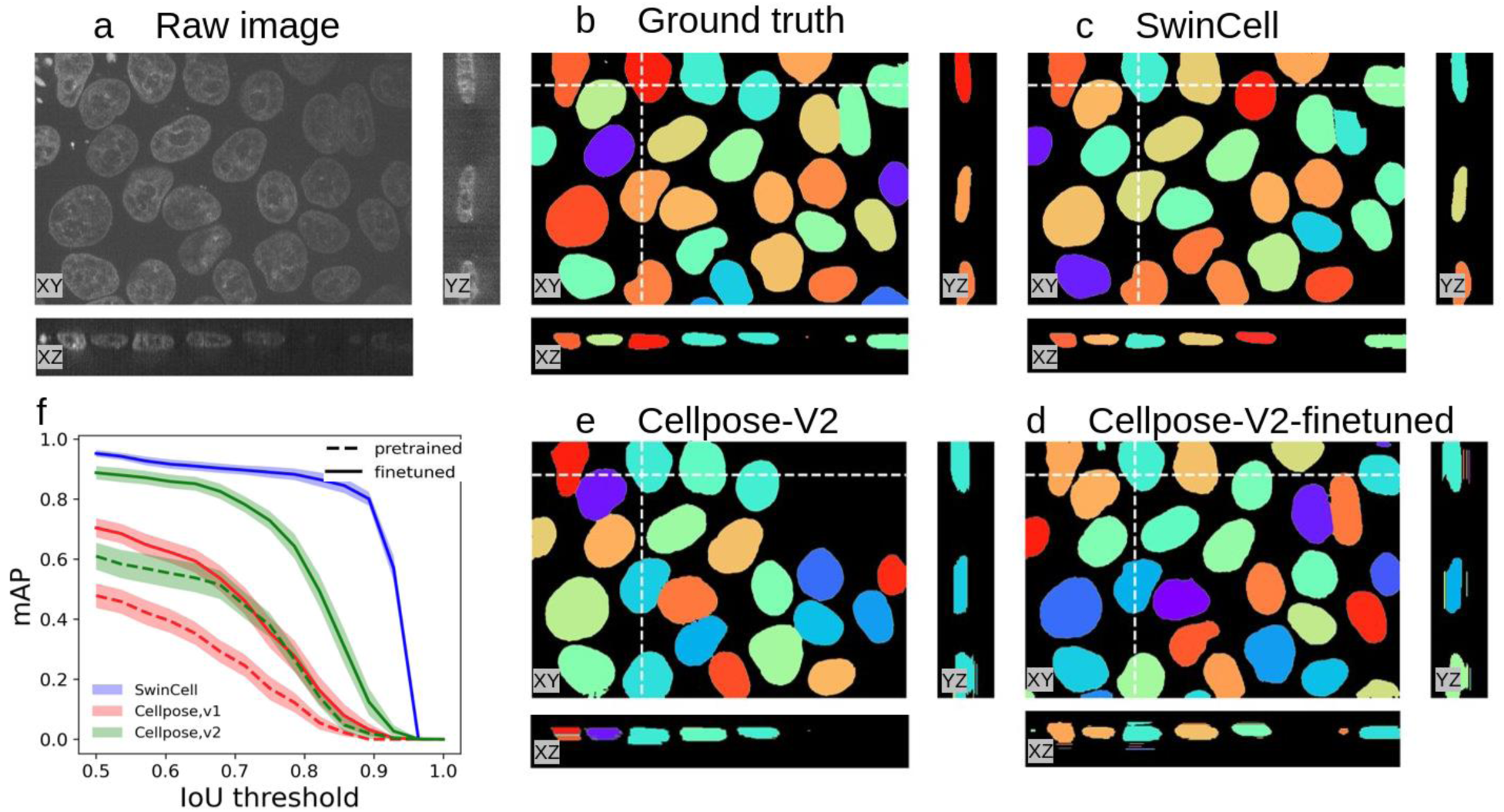
Segmentation performance of SwinCell on the Allen dataset. Representative slices of raw images (a), ground truth masks (b), Segmentation using SwinCell (c), Segmentation using pre-trained Cellpose model (d) segmentation using finetuned Cellpose model (e). The positions of the YZ, XZ slices are indicated by dash lines. **f**, Model performance measured by F1 score at different IoU (Intersection over Union) matching threshold. Two different Cellpose models were selected for comparison, and they are labeled as v1 and v2, respectively.

Four Cellpose variants (version 1, version 2, pre-trained with broad datasets, and fine-tuned with specific datasets) were used for comparison. The Cellpose model offers two methods for 3D segmentation using 2D-based models. One method predicts 2D flows and stitches them to form 3D flows for generating final 3D cell masks. The second approach converts predicted 2D flows into 2D masks, which are then stitched to form 3D masks. These methods are referred to as “Cellpose V1” and “Cellpose V2” respectively. Additionally, both the pre-trained ’Generalist’ models and the fine-tuned ’Specialist’ models were utilized for our comparative analysis (see the Methods section for more details).

Compared to the ground truth (**Fig. 2b**), SwinCell segmented all nuclei in 3D, both shapes and localizations with 0.83 recall, 0.97 precision, and 0.81 F1 score (**Fig. 2c**). As a comparison, Cellpose V1 and its fine-tuned version produced many false positives with many artifacts (**Extended Data** Fig. 2). Cellpose V2 produced fewer false positives but is incapable of predicting nuclei with weak fluorescence signals (**Fig. 2d**). Cellpose V2 fine-tuned outperforms V2 and can predict all nuclei, however, with a poor performance on predicting masks in the XZ and YZ planes (**Fig. 2e**).

We used mAP to comprehensively evaluate the performance of SwinCell and Cellpose variants across various Intersection over Union (IoU) thresholds, which served as matching criteria between ground truth and predicted cells, to determine true positives and false positives (**Fig. 2f**). Notably, the SwinCell model consistently outperforms Cellpose models across various IoU thresholds, demonstrating its superior precision and recall in cellular segmentation.

### Performance of SwinCell on a dense synthetic colon cell dataset

The colon dataset, featuring 3D digital phantoms of human colon tissue, effectively simulates both the physical and textural attributes of actual tissue structures. These intricately designed phantoms provide a representation of colon tissue to address the scarcity of labeled 3D datasets. The imaging dataset contains about 500 colon cells per image with a high cell density (**Fig. 3a**). The densely packed cells make the dataset suitable for testing the SwinCell performance.

**Fig. 3.**
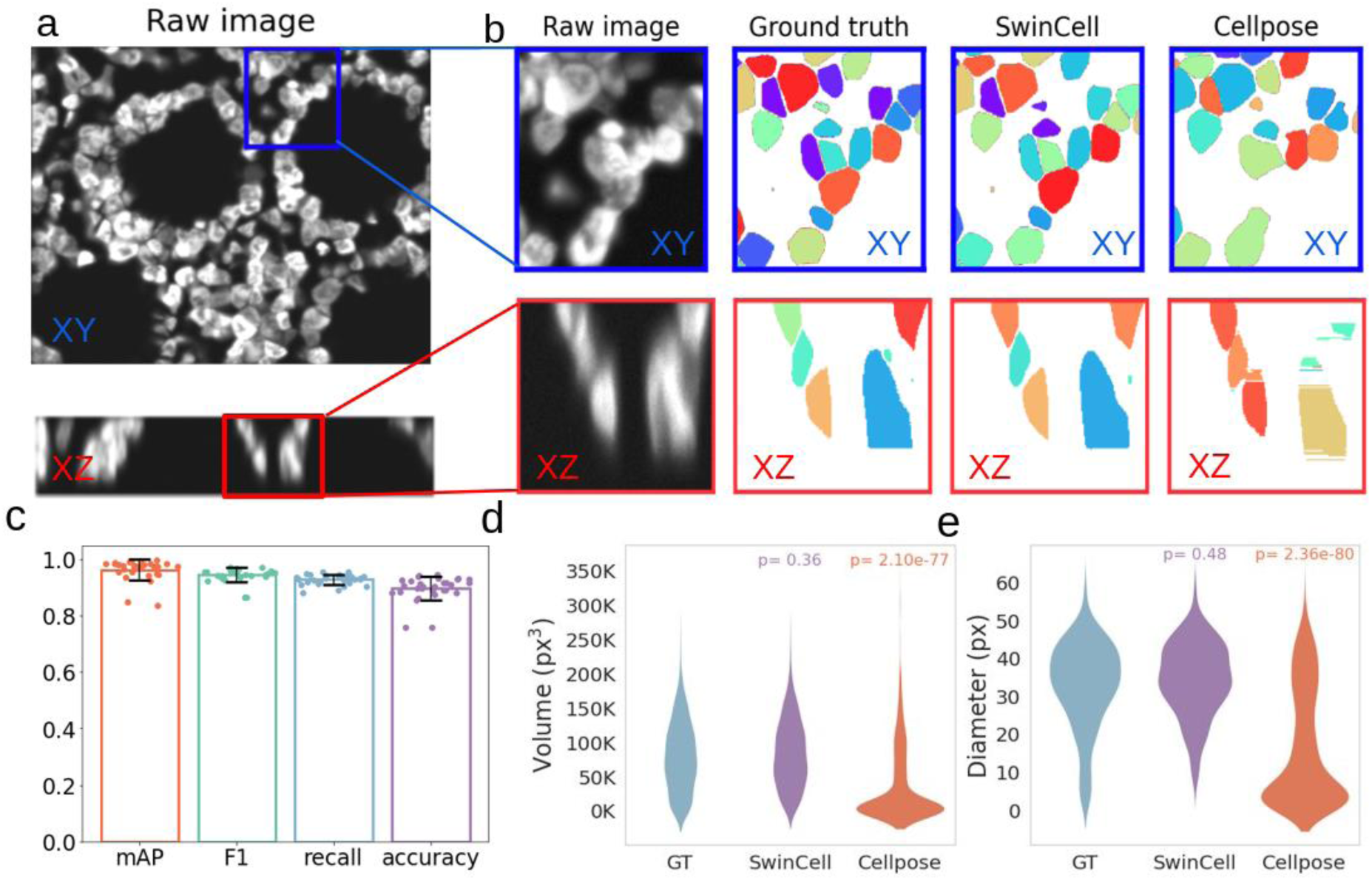
Segmentation performance of SwinCell on the colon dataset. **a**, Representative orthogonal slice and their zoomed-in views. Blue and red boxes highlight Regions of Interest (ROIs) in XY plane and XZ plane, respectively. **b**, Comparison of segmentation performance of SwinCell and Cellpose for the ROIs. Two orthogonal slice views of the segmented cell masks are shown. Different colors were used to define cell boundaries. **c**, SwinCell segmentation performance using Mean averaged precision (mAP), F1 score, recall, and accuracy. **d**, SwinCell segmentation performance against Cellpose using the volumes of segmented cells. GT, ground truth. The p-value on top of each violin plot shows a significant level of difference as compared with ground truth. Swincell predicted cell volumes and diameters are not significant from ground truth, while Cellpose predicted volumes and diameters are significantly different (p<0.005) from GT. **e**, SwinCell segmentation performance against Cellpose using the diameters of segmented cells as a metric. The width of each violin plot indicates the frequency of diameters within the dataset, with wider sections corresponding to higher number of cell masks with diameters around that value.

**Fig. 3a** exhibits the two orthogonal slices of a region of interest (ROI) extracted from the original 3D image. The corresponding ground truth masks, along with the segmentation results generated by SwinCell and Cellpose, are presented in **Fig. 3b**. The match between the SwinCell-segmented masks and the ground truth illustrates SwinCell’s capability to effectively delineate individual cells in both XY and XZ planes. In the sample image, SwinCell accurately segmented 427 cells out of 459 ground truth, with 32 false positives, yielding a precision of 0.93 and a recall of 0.93. This high accuracy is attributed to SwinCell’s 3D-based transformer model architecture, offering a broader perspective than traditional 2D models. In contrast, the Cellpose model produced 306 true positives and 751 false positives, reflecting a precision of 0.29 and a recall of 0.67. The higher rate of false positives with Cellpose is due to its 2D approach, which involves the slice-by-slice application of the 2D model to different XY planes and subsequent stitching slices to create a 3D volume, leading to fragmented segmentation in the XZ plane. This limitation is evident in **Fig. 3b**, where Cellpose shows efficacy in the XY plane but fragmented masks in the XZ plane. Different tunable parameters of the Cellpose model were tested to address this issue. However, the low precision (**Extended data Fig. 3**) persists despite varying stitching thresholds being applied.

We utilized the average values of mAP, F1 score, recall, and accuracy to evaluate the performance of SwinCell across the entire test dataset (**Fig. 3c**). All four metrics yielded mean value above 0.9, indicating high segmentation accuracy of dense cells within tissue images, thus establishing a solid basis for the post-segmentation cell analysis. To further compare the performance of SwinCell and Cellpose on the colon dataset, we focused on two features related to cell size, i.e., the cell diameter and cell volume. We calculated cell volume by the pixel count in 3D images, and estimated cell diameters as twice the maximum distance from a cell pixel to the cell boundary. **Fig. 3d,e** shows the comparison of the predicted cell volumes and cell diameters with the ground truth, respectively. Statistical analysis via Kolmogorov–Smirnov tests revealed that the distributions of SwinCell-predicted volumes and diameters were not significantly different from ground truth (p=0.36 for volumes and 0.48 for diameters), while the Cellpose-predicted volumes and diameters are statistically different from ground truth (p<0.005). These results show SwinCell’s capability to accurately predict 3D segmentation masks for densely interconnected cells, facilitating subsequent cellular analysis such as cell behavior, morphology, and biological process responses.

### End-to-end pipeline segmentation of in-house 3D cellular holotomography dataset

We extended our assessment of SwinCell’s applicability to diverse imaging modalities, incorporating in-house 3D holotomographic imaging data acquired using a Nanolive 3D microscope. Nanolive 3D images contain intricate cellular structures, including complex internal components such as nuclei and lipid droplets (**Fig. 4a**). These cellular details, while informative, pose segmentation challenges, influencing flow predictions and subsequent cell boundary delineation. Therefore, we employed Nanolive 3D dense cell images to demonstrate the end-to-end data processing and segmentation pipeline using SwinCell, from initial data annotation and model training to prediction.

**Fig. 4.**
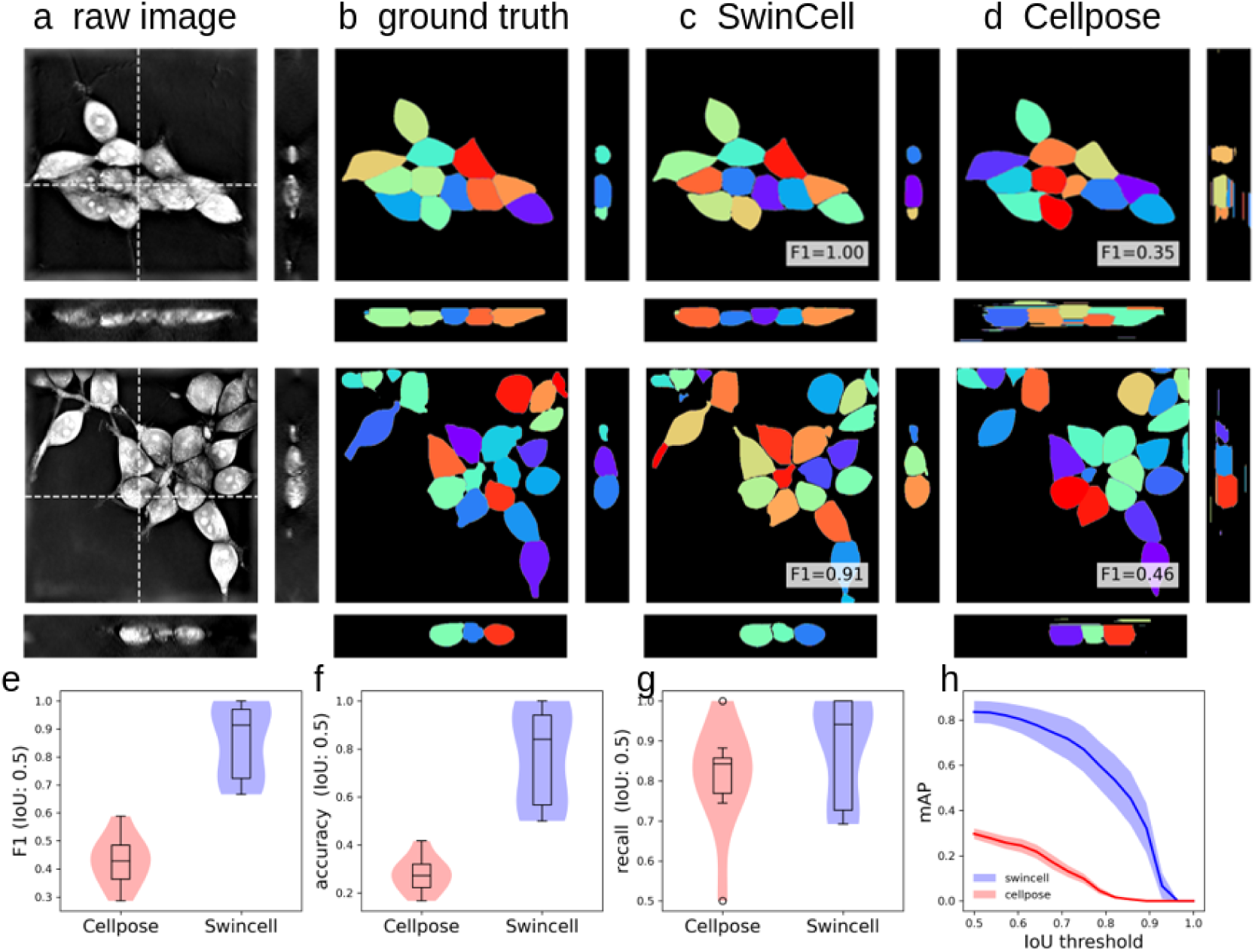
Segmentation performance of SwinCell on the in-house Nanolive dataset. **a**, representative views of two 3D cell-image slices of three orthogonal views. The positions of the slices are indicated by dash lines. **b**, Ground truth masks. **c**, Segmented cells masked by SwinCell. **d**, Segmented cells masked by Cellpose. The Matched F1 scores were displayed at the bottom right corner of each segmented image. **e-g**, Comparison of the segmentation performance of SwinCell (blue) against Cellose (red) using F1 score at 0.5 threshold (e), accuracy (f), and recall (g). **h**, Mean averaged precision (mAP) of SwinCell and Cellpose at different matching thresholds.

Our dataset comprises high-resolution 3D images of HEK293 adherent cells, with each image featuring approximately 20 densely packed cells. Two representative raw images with rich cellular features are shown in **Fig. 4a**. We manually labeled the raw images as 3D masks with examples shown in **Fig. 4b**. After training and prediction, the SwinCell model produces excellent segmentation results (**Fig. 4c,d)**, achieving an averaged precision of 0.84, recall of 0.88 and an F1 score of 0.86. As a comparison, the Cellpose model produces reasonable segmentation results on the XY plane but poorly segmented, i.e. fragmented, cells on the XZ and YZ planes. Overall, the Cellpose model achieves a lower precision, recall, and F1 score at 0.30, 0.81, and 0.43, respectively. These segmentation errors arise from challenges in seamlessly stitching different slices, a limitation inherent to the Cellpose model’s 2D-based approach.

The quantitative analysis through boxplots highlights the statistically better performance of SwinCell over Cellpose in terms of F1 score (**Fig. 4e**), accuracy (**Fig. 4f**), recall (**Fig. 4g**), and mAP (**Fig. 4h**). The mAP scores of SwinCell are higher than that of Cellpose across various IoU thresholds (0.5-1.0), providing further evidence of its superior 3D segmentation capability.

## Discussion

3D cellular tomography images by holotomography^1,2^, X-ray tomography^2^, confocal microscopy, cryo-electron tomography (cryo-ET)^4,23^, and other emerging imaging techniques are revolutionizing our observation and understanding of cellular structures and processes. Consequently, there is an urgent need for corresponding advances in image analysis algorithms to effectively analyze this data. Among these algorithms, the development of advanced 3D cell segmentation algorithms is pivotal. In this work, we address this need by developing the SwinCell model that leverages the innovative Swin-transformer^6,19,24^ model architecture, which has shown remarkable proficiency^6,25,26^ in handling complex 3D image data by effectively capturing both local and global contextual information in multiple scales. In addition, SwinCell integrates the cell flow gradient prediction^10,14^ algorithm with Swin-transformer, enabling the model to effectively distinguish individual cells within dense clusters from multiple modal data of confocal microscopy of nuclei, colon tissue cells, and densely cultured cells. The utility of SwinCell to various 3D data types suggests its broad application in studying cellular events.

Prior to this work, substantial efforts were devoted to applying 2D deep-learning models, for example, U-net and Mask-RCNN, to segment 2D slices of 3D tomograms. These models essentially segment 2D slices of a 3D tomogram and stitch pieces together for a 3D segmentation. Representative examples are Cellpose^10,11^ and Mesmer^9^. These models work well on 2D cellular images. Nevertheless, they showed limited performance in segmenting 3D cellular images, since the predicted 2D masks lack consistency across slices. Notably, throughout both the training and inference phases, the model operates under the assumption that cellular diameters remain consistent across different slices. This assumption, however, does not hold in reality. The cells usually have a spherical shape in 3D volume resulting in variations in size when they are intercepted at different axis locations. Consequently, the models’ performance is adversely affected due to the misestimation of cell diameters. In contrast, the SwinCell model has a more integrated view of the cell shapes and thus is better at segmenting the cells’ volumes. Even for the best performer of Cellpose with extensive fine-tuning, SwinCell shows superior performance on all matrices used for the comparison (**Figs. 2-4**).

Swin-transformer uses shifted windows multi-head self-attention (SW-MSA)^19,24,27^ mechanism that allows for the model to transmit the learned attention information between windows (**Fig. 1**), thus giving the model a more integrated view of the whole cellular image. The Swin-transformer is in theory more capable of predicting a smooth flow than conventional U-Net architecture. By combining the in-depth 3D analysis with the efficiency of the 3D Swin-transformer, our model offers a novel solution for accurate 3D segmentation. To better understand its performance versus U-Net, we performed an ablation study (**Extended Data** Fig. 4) and found that SwinCell predicted smoother flows as compared with the U-Net model. The predicted flows are more consistent with the ground truth. The flow plays an important role in the proposed instance segmentation algorithm, as a result, the predicted cell masks showed higher prediction accuracy (F1 score) as compared with U-Net algorithm. **Extended Data** Fig. 4i-l shows a systematic comparison of the performance of U-Net without attention mechanism and SwinCell in terms of F1 score, accuracy, and recall. SwinCell demonstrated a higher F1 score and accuracy when tested, despite there being no significant difference in the recall values. Through the comparison, we show that SwinCell can predict smother flows and thus more accurate segmentation masks. In contrast, the flows predicted by U-Net usually show discontinuity and lead to false positive predictions. As shown in **Extended Data** Fig. 4l, the false positive predicted by U-Net is significantly higher than that of SwinCell.

Computational constraints pose a significant challenge for complex 3D model architectures. 3D model architectures require substantial computational resources in terms of CPU or GPU memory. Addressing this challenge necessitates innovative approaches in model design. By focusing on smaller, local windows instead of the entire input image, Swin-transformer significantly reduces the complexity of the self-attention mechanism. Traditional transformers exhibit a computational complexity of O(N^2^) with respect to the input image size N, whereas the Swin-transformer reduces this to O(N) for each window, resulting in lower overall computational demands^24^. In addition, the shift windows mechanism in the Swin-transformer facilitates cellular segmentation in 3D. The neighboring patches from the input image allow the model to consider information from adjacent patches, which helps reduce the boundary artifacts and improve continuity across patch boundaries. The Swin-transformer architecture used in SwinCell is thus less sensitive to the internal textures of the cells and less sensitive to patch size limitations for flow prediction and subsequent segmentation. In addition, the Swin-transformer architecture predicts smoother gradient flows along each dimension of the images. Finally, the Swin-transformer can achieve good performance with fewer parameters compared to traditional attention-based models^6,26^, leading to better generalization^28^.

One key goal of cellular segmentation is the automation of quantitative analysis of the cells. It is therefore important to ensure that the segmented cell shapes and volumes are consistent with the ground truth. As shown in **Fig. 2**, the volumes and diameters of SwinCell-predicted cells showed no significant difference with ground truth labels, providing reliable measurements of the cellular morphology and behavior. To test the feasibility of the SwinCell model in different scenarios, we applied it to different datasets of tissue and single-cell images, nucleus, fluorescent images as well as cultured cell images. Most of the cellular images contain dense cells connecting each other, making the segmentation work challenging. SwinCell showed good performance in all the datasets, demonstrating its capability of multimodal segmentation for various cellular quantification and analysis approaches.

One challenge of 3D segmentation is the availability of labeled 3D datasets. While we show that SwinCell can segment various types of 3D cellular images, the development of a universally applicable 3D “generalist” model remains a challenge, primarily due to the scarcity of labeled 3D cellular data. Creating 3D segmentation masks for training data is a labor-intensive and time- consuming task^17^, often resulting in a shortage of high-quality training data for 3D models. In this study, we addressed this issue by employing synthetic data^29^ generation techniques, which can create vast amounts of artificial data with accurately labeled ground truth, thus mitigating the training data shortage. Additionally, we utilized human-in-the-loop methodologies^9,11^ to enhance and expedite the quality improvement of our Nanolive dataset, whereby human annotators iteratively refine the model’s predictions, thereby improving training data quality.

Although not applied in the current study, self-supervised learning^27^ represents a promising direction for future work. The architecture of the Swin-transformer, with its inherent flexibility and capacity for capturing complex patterns in data, is particularly well-suited for self-supervised learning approaches. By leveraging extensive amounts of unlabeled data, the Swin-transformer can be pre-trained to develop an intrinsic understanding of 3D image features, which can later be fine-tuned with a smaller set of labeled data. As the repository of available 3D cellular datasets expands, we are optimistic about employing the SwinCell architecture for the development of a comprehensive, universally adaptable 3D segmentation model.

## Methods

The SwinCell algorithm is developed using Python 3.8^30^, Pytorch 1.11.0^31^, Jupyter-notebook^32^, Matplotlib^33^, and Seaborn^34^ are used to generate the figures in the paper.

The 3D SwinCell algorithm was tested with two public datasets and one in-house dataset. These three datasets contain cellular images or nucleus images in different scenarios.

### Public Datasets

Colon tissue dataset^29^ is a challenging dataset with densely connected cells of the colon. It includes 30 synthetic images of human colon tissue. As described in reference^29^ the dataset was generated using the virtual microscope imitating the microscope Zeiss S100. The images are provided in various formats and the 3D-TIFF format was used in this paper. The ground truth semantic segmentation labels are also provided along with this dataset. As described in the model training section, the semantic labels are transformed to instance labels before model training, with each cell assigned a unique integer value to separate individual cells.

The Allencell dataset is provided from reference^22^. In this study, a total of 100 3D images were utilized, and distributed across training, validation, and testing phases. The dataset was split with 60% of the images allocated for training, 20% for validation, and the remaining 20% used for testing. This distribution was chosen to ensure a comprehensive learning process while providing adequate data for the validation and performance evaluation of the model. The dataset includes fluorescent protein marker channels that are used for identifying different organelles in the cell. The corresponding segmentation labels are also provided for each channel in the dataset. In this work, the DNA cell nucleus channel was used to test and evaluate the SwinCell algorithm.

### Holotomography cellular image collection

As described previously^2^, adherent HEK293 cell (ATCC Cat# CRL-1573) was cultured in Dulbecco’s Modified 305 Eagle’s Medium (DMEM) supplemented by 10% (v/v) fetal bovine serum and 100 units/ml 306 of penicillin-streptomycin (Cat# 15140122, Thermo Fisher). The dish was placed in a stage-CO2 incubator on a 3D Cell Explorer-fluo light microscope (Nanolive). The incubator was operated at 37°C, 5% CO2, and 100% humidity. Image data were collected at different locations and the resultant images were saved in 3D-TIFF format.

### Data preparation and annotation

For this study, all input data was transformed and saved in 3D-TIFF format for easier data manipulation. To accommodate the varying intensity distributions present in the input images, a normalization procedure was applied to each image. The 1st and 99th percentiles of the intensity values of the input image were adjusted to 0 and 1, respectively. Such normalization ensures consistent intensity scales across different images, allowing the model to process and learn from the data more effectively.

The collected 3D cellular Nanolive data was manually labeled with ITKsnap^35^ (www.itksnap.org). First, a binary mask was manually annotated, designating the cellular regions as 1 and the background regions as 0. To enhance the smoothness of the labeled mask, binary image opening and hole-filling operations were applied. Subsequently, a Python script was employed to transform the binary mask into an instance mask, designating the background as 0 and assigning a distinct positive integer to each cell. Then the resultant labels are manually fixed to ensure that the cells are distinguishable. In cases of densely populated cell regions, several iterations of manual corrections and smoothings were executed to enhance the label accuracy.

### Model architecture

The 3D variant of the Swin-transformer^19,24^ is used in this study. The model adopts an encoder- decoder architecture^12,19^. The model has 3 stages, and the feature representation of each stage is decreased by a factor of 2 after the patch merging layer. Unlike the conventional U-Net model^12^, the Swin Transformer incorporates self-attention modules within its encoder part. The model incorporates the flow prediction in Cellpose into a Swin-transformer backbone, effectively transforming it into an instance segmentation model. The model predicts four channel outputs simultaneously: a cell probability map and three flow maps along the XYZ directions. Following the Cellpose segmentation pipeline, the predicted XYZ flow maps are subsequently used to group individual cells with the gradient tracking method described in the method section. After gradient tracking, each cell instance will be labeled with a unique integer ID. The model was constructed using Pytorch^31^, cuda 12.0 and MONAi^36^. Numpy^37^, and Scipy^38^ libraries are used for basic matrix and image manipulation.

### Flow Generation

The flow calculation process mimics the process of heat diffusion and is adopted from the Cellpose algorithm^10^. Briefly, a flow source was introduced at the center of each cell in the ground truth masks, and spatial derivatives were calculated along the specified direction. Finally, gradient values were assigned from one boundary to the boundary of the other side. The flow values were normalized to [-2, 2] before model training.

The same flow generation process was applied for all the datasets tested in this work. Some of the datasets (e.g. the colon dataset) only contain semantic labels, where the cell regions are labeled as 1 and background regions are labeled with 0. For easier flow simulation, the semantic labels were translated to instance labels where each cell is given a different value (Fig. 1b). The Python script provided in this work can be used to efficiently translate labels into instance segmentation labels in the data preparation stage, with each cell labeled with a unique integer value.

### Model training

The model input size was set to (128,128,32) and the ROIs were randomly generated from the input images and ground truth labels. In this work, the Monai^36^ package was used to efficiently generate image patches of a fixed ROI size randomly from the input images. A feature size of 48 was used as the embedding size. We trained the model with the Adam optimizer in Pytorch and the initial learning rate was set to 1e-4 with a 1e-5 decay for 10K iterations. All training was performed on two NVIDIA RTX 3090 (24 GB) GPUs, and the batch size was set to 1 per GPU.

We use the sum of L2 loss the flows and cross-entropy of the cell probability as the loss function:

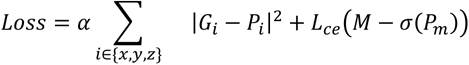

where α is the hyper-parameter to adjust the relative importance of flow loss and cell probability loss. We used α=5 according to the Cellpose paper^10^. *G_x_*, *G_y_* and *G_z_* are the 3D flow volumes of ground truth, where each voxel indicates the flow of that voxel in the *x*, *y* and *z* direction, respectively. Likewise, *P_x,_ P_y_* and *P_z_* represent the corresponding predicted 3D volumes. *M* is the 3D probability mask of the ground truth, while *P_m_* is the predicted probability mask. *L_ce_* is the standard binary cross-entropy loss and *σ* is the sigmoid function.

### Model evaluation

The model evaluation process implemented a 5-fold cross-validation strategy, with 60%, 20% and 20% images used for training, evaluation and testing, respectively. For a fair comparison, the same data splits were applied to train the SwinCell model and all other models in this paper.

Different versions of the Cellpose model are used to compare the model performance. The Cellpose algorithm utilizes 2D-based U-Net style models and thus can only make 2D predictions. However, this algorithm provided two pipelines to deal with 3D cellular data with 2D models. The first method involves predicting 2D flows, stitching the 2D flows to 3D flows, and then producing the final 3D cell masks. The other method is to convert the predicted 2D flows to 2D masks, and then stitch the 2D masks to 3D masks. Both methods are evaluated with the Allencell dataset in this paper, and they are referred to as “Cellpose V1” and “Cellpose V2” respectively (Figure 2). As shown by the mean average precision (mAP) values at different thresholds in Fig. 2d, the second algorithm (v2) consistently shows better results than v1.

The Cellpose model is typically known as the ’Generalist’ model, renowned for its proficiency across various cell types and imaging modalities. For tasks that require more specificity, a ’Specialist’ model can be fine-tuned from this Generalist model, tailored to specific cell types or imaging methods for enhanced performance. In our study, we used both the pre-trained Generalist and the fine-tuned Specialist versions of the Cellpose model for a comprehensive comparative analysis.

The pre-trained Cellpose models are downloaded from the Cellpose website without modification. For the finetuned models, model training was performed with the same data splits as the SwinCell model.

A few evaluation metrics were used to measure the accuracy of cell localization and pixel classification. The number of correctly segmented cells is estimated using various Intersection over Union (IoU) matching thresholds. A predicted mask is labeled as a true positive if its IoU with a ground truth label exceeds the specified threshold. Precision is calculated as True Positives / (True Positives + False Positives) and recall is calculated as True Positives / (True Positives + False Negatives). Mean Average Precision (mAP) was calculated as the average value of AP (Average Precision) values across the test subset of images. The F1 score of the predictions is calculated as the harmonic mean of precision and recall. This score provides a single metric that balances both precision and recall. The F1 score is particularly useful in scenarios where an uneven class distribution might exist, as it maintains a balance between the model’s sensitivity and its ability to correctly identify negatives. By utilizing both mAP and F1 scores, we ensure a comprehensive evaluation of SwinCell’s performance in accurately segmenting and identifying cells within varied and complex 3D imaging environments.

## Acknowledgments

This work was supported by Laboratory Directed Research and Development Program LDRD21-013 and the U.S. Department of Energy (DOE), Office of Biological and Environmental Research (KP1601011).

## Author contributions

X. Z, Y. L, and Q.L. designed the study and experiments. X. Z. and Z. L. performed the experiments. All authors analyzed the data. X. Z. and Q.L. wrote the manuscript with help from other coauthors.

## Competing interests

Authors declare no competing interests.

Correspondence and requests for materials should be addressed to X. Z., Y. L., or Q.L.

## Data availability

The 3-dimensional cellular data using a Nanolive homography microscope are available upon reasonable request.

The Allencell (hiPS) dataset is available at: https://www.allencell.org/data-downloading.html. A tutorial for downloading the dataset is available at https://github.com/AllenCell/quilt-data-access-tutorials/tree/main

The colon dataset is available from https://cbia.fi.muni.cz/datasets/

## Code availability

The codes are freely available at https://github.com/xzhang0123/SwinCell

---

**Extended Data Fig. 1.**
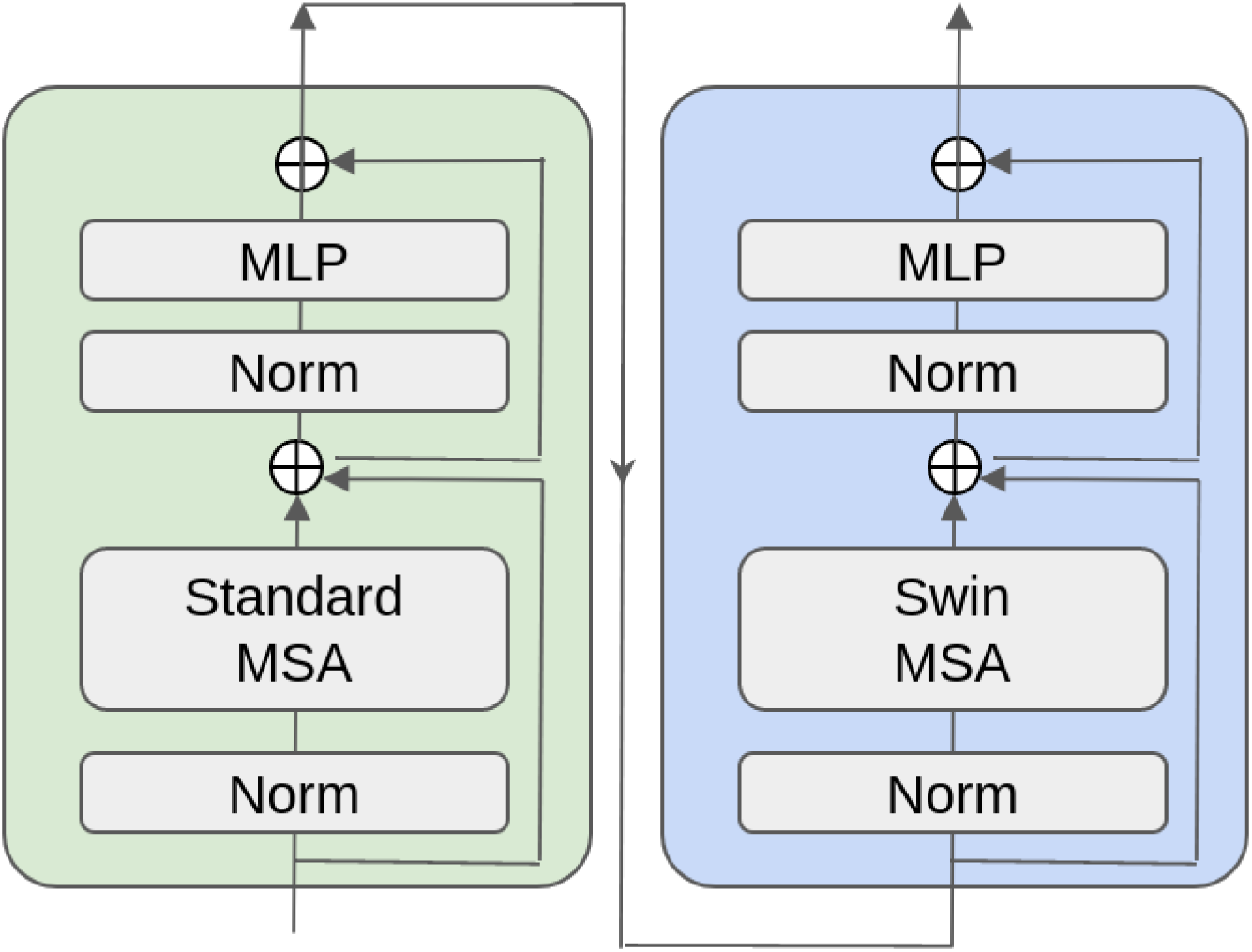
A Swin unit consists of two consecutive self-attention modules. The first is a standard Multi-head Self-Attention (MSA) module and the second is a Swin Multi-head Self-Attention (Swin-MSA) module. MLP denotes Multi-Layer Perceptron.

**Extended Data Fig. 2.**
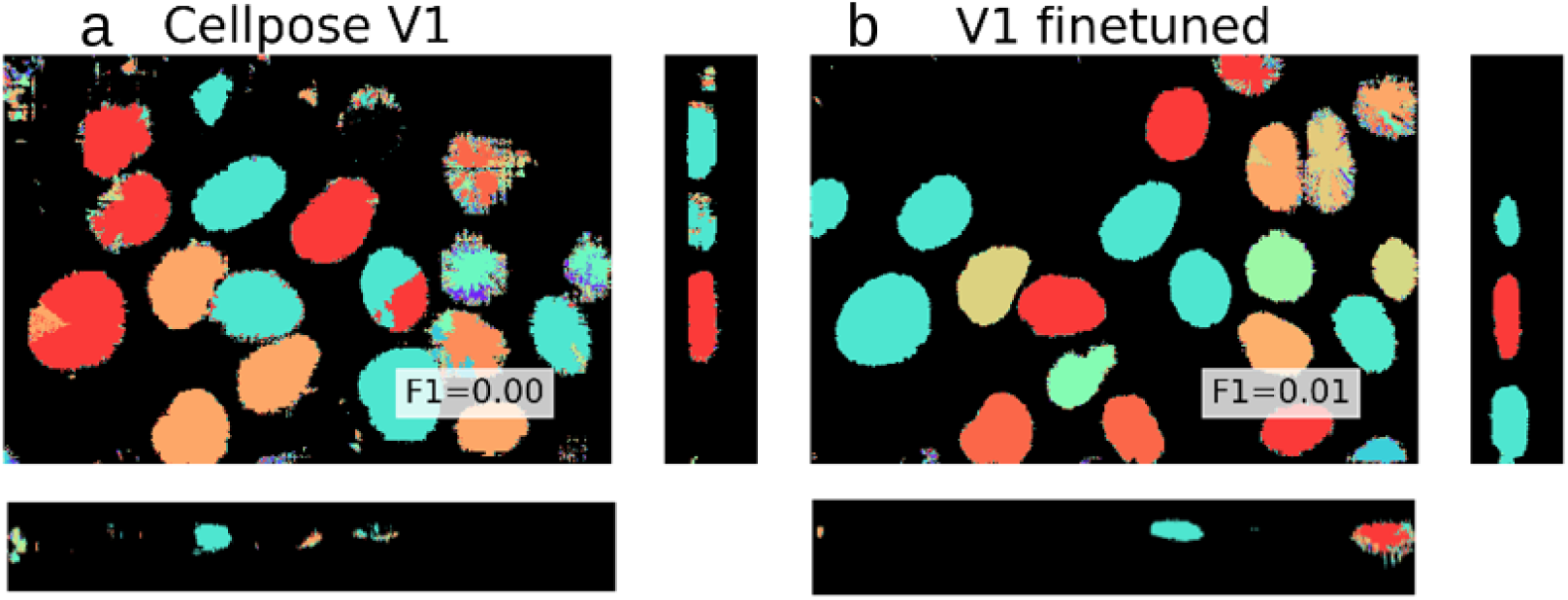
Representative segmentation results of Cellpose models on the Allencell dataset with the first approach (v1). This method involves predicting 2D flows, stitching them into 3D flows, and subsequently generating the final 3D masks. Further details are provided in the Results section. **a**. Segmentation using the pre-trained Cellpose model. b. Segmentation using finetuned Cellpose model. Matched F1 scores of the corresponding images are displayed at the bottom right of the image.

**Extended Data Fig. 3.**
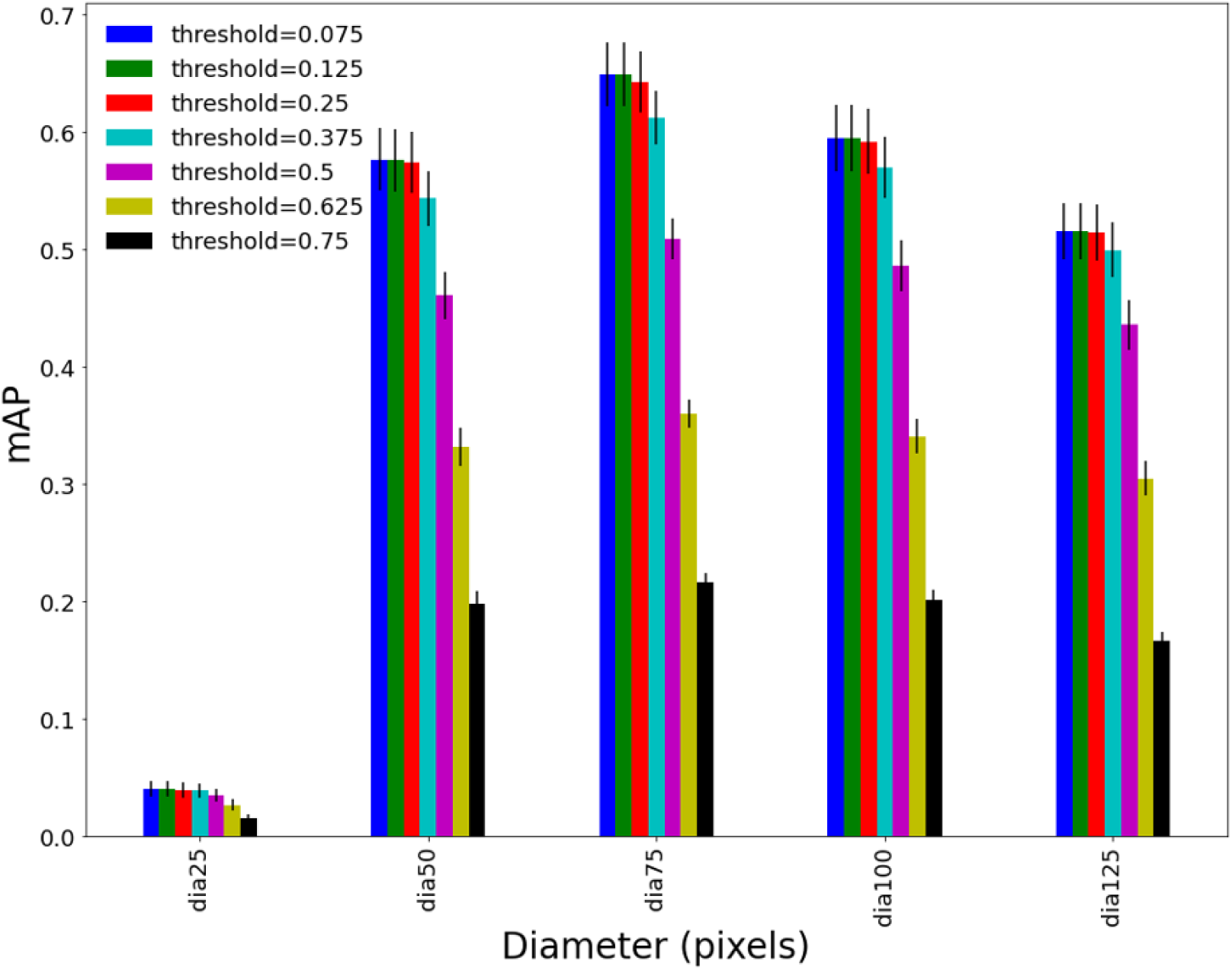
Model performance of Cellpose using different parameters. Two parameters were optimized for the Cellpose model: cell diameter and matching threshold for stitching different slices of the volume. Various combinations of these two parameters were tested, and the finetuned Cellpose model with the best performance was selected to compare with the performance of the SwinCell model.

**Extended Data Fig. 4.**
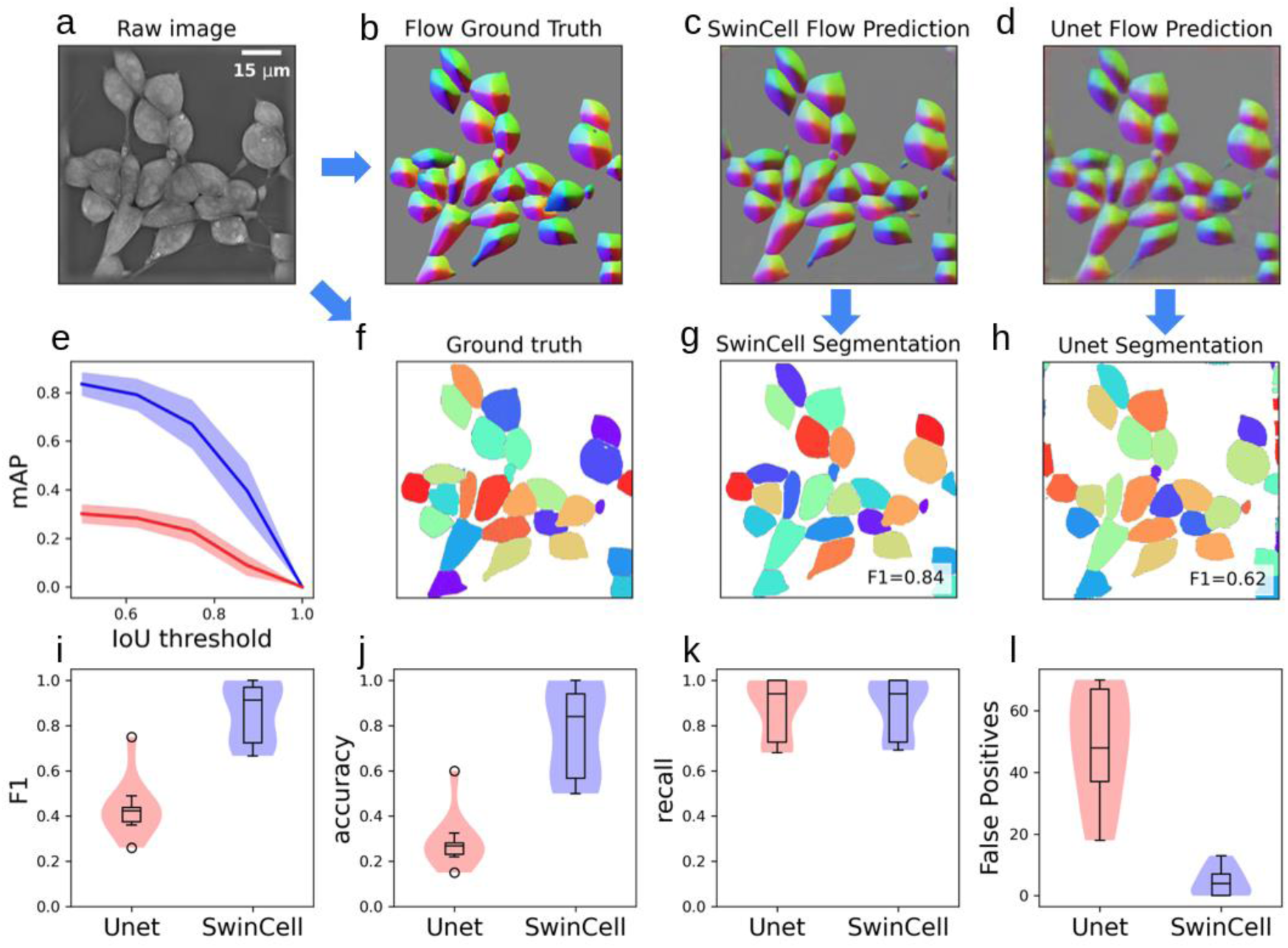
Ablation analysis on the model performance of SwinCell model with the corresponding Unet model. **a**, sample Nanolive image to be segmented. **b**, Ground truth flow generated from ground truth cell instance labels. **c-d**, Model predicted flow by SwinCell and Unet, respectively. **e**, Mean averaged precision (mAP) of SwinCell (blue) and Unet (red). **f-g**, comparison of ground truth (f), SwinCell (g) and Unet (h) predicted segmentation masks. The Matched F1 scores are displayed at the bottom right corner of each segmented image. **i-l**, Additional metrices comparing Unet (red) and SwinCell (blue) of F1 score (i), accuracy (j), recall (k) and False Positive (Number of false predicted cells) (l).

## Notes

### Competing Interest Statement

The authors have declared no competing interest.

